# Disentangling RNA evolution and thermodynamics in genomic language models

**DOI:** 10.64898/2026.05.28.728275

**Authors:** Yuchen Xu, Nevin N. Pai, Hannah K. Wayment-Steele

## Abstract

Genomic language models (gLMs) trained only on large-scale nucleic acid sequence data seem to capture signals of RNA structure, yet the specifics of how remain unclear. Using the categorical Jacobian (CJ) operation, a model-agnostic operation for querying pairwise dependencies, we systematically compared three flagship gLMs: RNA-FM, Evo 2, and gLM2. We found that CJ signals recover base pairs supported by evolutionary covariation analyses, consistent with findings in protein language models. Surprisingly, CJ also recovers base pairs lacking evolutionary support but predicted by biophysical nearest-neighbor models. Is it possible gLMs have “learned” RNA thermodynamics? We noticed nearest-neighbor RNA folding models often predict reflected structures when given reversed sequences, consistent with these models’ modular and grammar-like nature. We leveraged this observation to create a simple “mirror test” that we found gLMs routinely fail, indicating they have not learned generalizable biophysics-based rules for RNA structure. Nevertheless, their apparent thermodynamic signal potentially confounds interpreting gLM pairwise dependencies as evidence of evolutionary conservation. We therefore introduce a method using synthetic sequences as a control for detecting significant learned signal. Our results demonstrate that gLMs can mimic thermodynamics through learned sequence context rather than general physical principles, but solutions exist for disentangling patterns in language models.

## Introduction

RNA molecules play central roles in cellular processes, with their functions often closely linked to their secondary and tertiary structures^1,2^. Understanding how RNA structure arises from sequence is therefore a fundamental problem in molecular biology. Over the past few decades, biophysical models based on thermodynamic principles have been developed to predict RNA secondary structure from sequence, including widely used methods such as ViennaRNA and related “nearest-neighbor” models^3,4^.

Recently, the emergence of genomic language models (gLMs) has opened potentially unprecedented routes for structure modelling and discovering novel structure and function of RNAs. These models are trained on large sequence repositories and then finetuned for a variety of structure prediction tasks^5–7^. Their success in these tasks has led to growing interpretation that gLMs encode internal representations of RNA structural rules^8–10^. Similar interpretations have previously emerged in protein language models, where pairwise sequence dependencies learned from sequence are associated with structural contacts and evolutionary couplings. In protein structure modelling, evolutionary covariation has long served as a powerful proxy for structural conservation by leveraging correlated mutations between positions that preserve structural interactions^11–14^.

However, the interpretation of evolutionary covariation in RNA is often fraught. For example, structural conservation in long noncoding RNAs such as MEG3 and HOTAIR has been proposed in some studies but questioned in others, in part due to limitations in covariation-based inference and statistical interpretation^15,16^. Such methods, such as R-scape^17,18^, rely on detecting evolutionary covariation from multiple sequence alignments (MSAs). However, the statistical power of these approaches depends heavily on alignment depth, sequence diversity, and alignment quality, which vary substantially across RNA families^19^. More robust deep- learning-guided frameworks for detecting evolutionary conservation of structure in RNAs would be transformative, and gLMs have emerged as an appealing route toward this^20^.

Recent work has further suggested that apparent RNA structural signals may emerge from relatively simple sequence statistics^21,22^. Compact stochastic context-free grammar (SCFG) models trained without structural annotation were shown to recover canonical RNA base-pairing rules and achieve competitive secondary-structure prediction performance. These findings raise the possibility that gLMs may similarly recover structure-associated signals through large-scale extraction of sequence statistics, potentially approximating simple biophysical base-pairing relationships. If so, model-derived pairwise signals would be expected to exhibit relatively uniform enrichment for base pairs across RNA families and obeying rules of the grammar.

In this study, we apply the categorical Jacobian (CJ) operation^23^ to systematically dissect the pairwise signals of several gLMs across diverse RNA datasets, including models trained on mixed sequence modalities (e.g., gLM2^24^ and Evo2^25^) and RNA-specific models (e.g., RNA-FM^7^). Rather than interpreting a specific internal mechanism such as attention, CJ measures how perturbations to the input sequence influence model outputs, capturing full input-output dependencies through the entire model. As a result, CJ is architecture-agnostic, enabling direct comparison across models with substantially different internal designs, including transformers (RNA-FM, gLM2) and the hybrid attention/convolution stack used by Evo 2. This property addresses well-recognized concerns that attention weights alone are not always faithful explanations of model behavior^26^.

When first introduced, the CJ operation demonstrated that the protein language model ESM-2^27^ encodes protein evolutionary couplings as a direct result of masked language modelling, prior to further supervision on contacts^23^. More recent studies have applied CJ to gLMs such as gLM2, Evo^9^ and Evo 2 to provide individual examples of where these models encode RNA structure-associated signals. Here, we compare CJ signals to two established and contrasting types of RNA pairing evidence: evolutionary covariation identified by R-scape^17^, and biophysical pairing propensities predicted by nearest-neighbor thermodynamic models, EternaFold^28^ and ViennaRNA^3^. R-scape detects statistically significant covarying base pairs from sequence alignments and is widely used as evidence for conserved RNA structure, whereas nearest-neighbor models estimate RNA base-pairing energetics from experimentally derived thermodynamic parameters. We observe that gLM CJ signals predict some base pairs with evolutionary covariation support, consistent with prior observations in proteins. However, CJ signals predict base pairs that lack detectable covariation but are supported by biophysics-based “nearest-neighbor” models.

We devised a simple yet incisive experiment to test if CJ signals might encode grammar akin to nearest-neighbor models. We observed that nearest-neighbor models, if given a reversed sequence, will often predict a structure “mirrored”, i.e. inverted, from the structure of the original sequence. By use of this “mirror test”, we found that CJ signals are ablated when RNA sequences are reversed. This observation challenges the idea that gLMs recover fully generalizable grammatical rules of RNA pairing, which would likely be a prerequisite for transferable thermodynamic or biophysical representations. Together, these findings point to a more complex relationship between gLM pairwise signals, evolutionary covariation, and thermodynamic pairing propensity.

This complexity creates an important challenge for interpreting gLM pairwise signals: if CJ signals can mimic biophysics-based base-pairing probabilities, CJ signal at a proposed base pair cannot be directly interpreted as evolutionary conservation. Finally, we propose a significance test that compares CJ signal from natural RNAs to that from synthetic RNAs designed to fold into the same structure. Positions where natural sequence CJ signal exceeds the distribution of signals from synthetic sequences are candidates for evolutionary inspired interactions from CJ signals. This framework offers a practical path forward for using gLM-derived pairwise signals to evaluate patterns of evolutionary conservation.

## Results

### Categorical Jacobian base pair prediction varies widely by RNA family

We used the categorical Jacobian (CJ) operation to assess the degree to which pairwise information learned in language models reflects RNA secondary structure. In brief, the CJ operation performed on an RNA sequence *x* of length *L* involves systematically mutating each position and measuring the resulting changes in model outputs at each other position (Fig. 1A). Let *f_j_*(*x*)*_b_* denote the model logit at position *j* with nucleotide *b*. For each position*i*, we generate perturbed sequences *x*^(*i→a*)^ by substituting the nucleotide at position *i* with nucleotide *a* ∈ {*A, C, G, U*}, and compute:

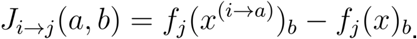

**Figure 1.**
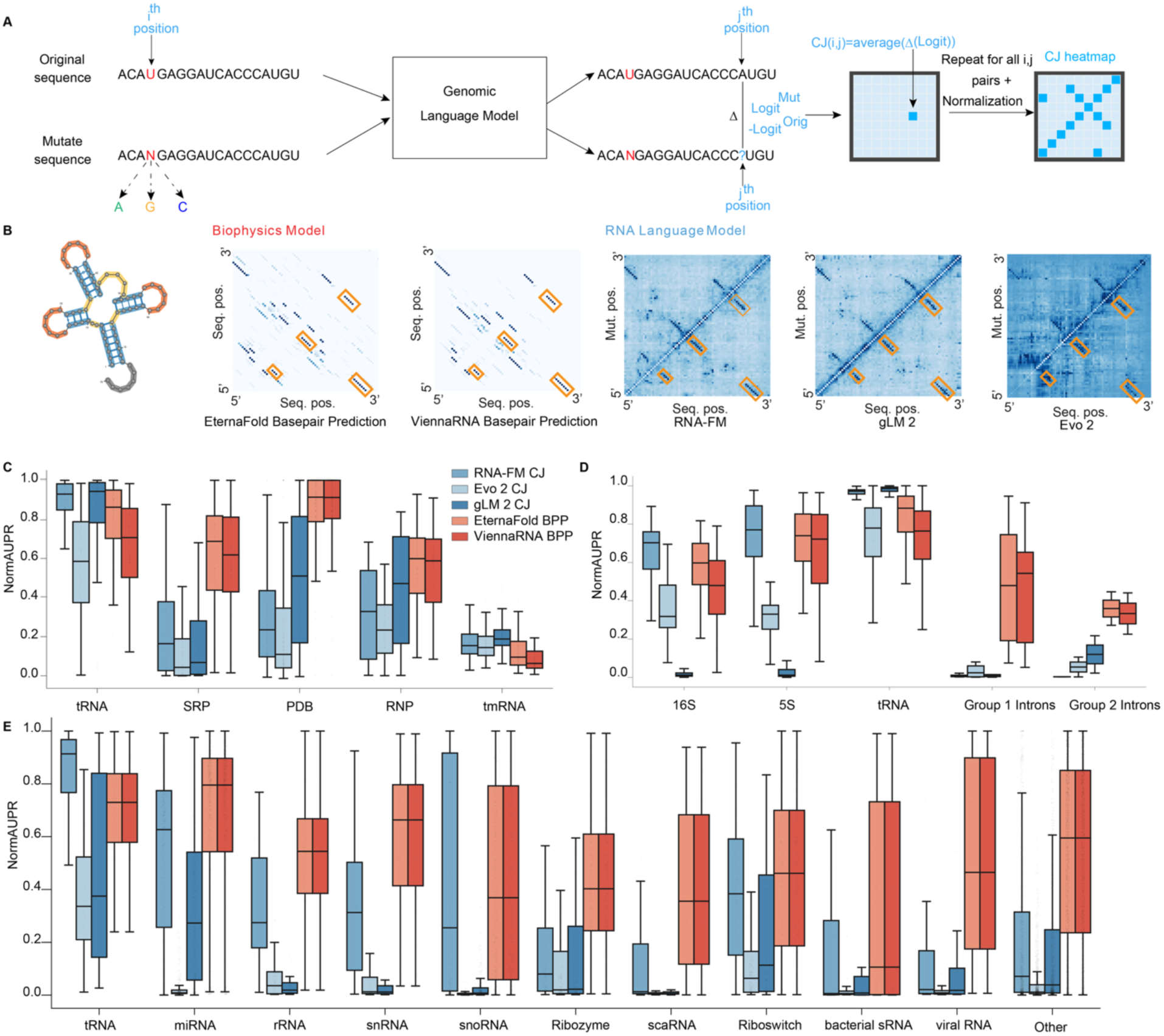
Categorical Jacobian signals vary widely in predictive power for base-pairing in noncoding RNAs across RNA families. **(a)** Schematic illustration of the categorical Jacobian (CJ) operation used to quantify model-derived pairwise dependencies between nucleotide positions from genomic language models (gLMs). **(b)** Representative example for a tRNA (bpRNA_CRW_30836), showing annotated secondary structure, base-pairing probability (BPP) maps from two biophysical models (EternaFold and ViennaRNA), and CJ interaction maps from three gLMs (RNA-FM, gLM2, and Evo 2). Nucleotide pairs corresponding to annotated base pairs are highlighted by yellow squares on the CJ and BPP heatmaps, demonstrating that in this example, the CJ signals capture known tRNA stems. Distribution of normalized area under the precision–recall curve (NormAUPR) for different models across **(c)** bpRNA datasets, **(d)** stratified by RNA family within the CRW dataset from bpRNA, **(e)** stratified by RNA family within the Rfam dataset, showing substantial variation in CJ performance across RNA families relative to biophysical models.

This yields a four-dimensional tensor *J* ∈ ℝ^*L*×4×*L*×4^, representing the sensitivity of outputs at position *j* to perturbations at position *i*. After mean-centering across tensor dimensions, we computed pairwise interaction scores using the Frobenius norm, symmetrized the resulting matrix, set diagonal entries to zero, and applied average product correction (APC) to reduce global background biases.

Biophysical models that have been under development for decades have predictive power for tRNA structure: this is illustrated by comparing the structure of glutamine tRNA from *Leptospira interrogans* (bpRNA_CRW_30836)^29^ to base-pairing probability (BPP) maps from two models, EternaFold and ViennaRNA (Fig. 1B). Notably, the CJ-derived signals from three nucleotide language models — RNA-FM, Evo 2, and gLM2 — all show features consistent with known tRNA stems (orange boxes). We analyzed the CJ signals for these three gLMs across 1,250 RNA sequences from six datasets derived from bpRNA_1m_90^29^. For each sequence, we computed a normalized area under the precision–recall curve (NormAUPR), which quantifies how strongly top-ranked CJ pairs are enriched among annotated base pairs (see Methods). For comparison, we performed the same analysis using base-pairing probabilities from EternaFold and ViennaRNA. We compared to EternaFold given its state-of-the-art predictive power for observables based on base-pair probabilities in ref. 28. We also compared to ViennaRNA as a more widely-used method; we ran ViennaRNA at 60°C, which was shown to have higher predictive power than its default temperature for ensemble-averaged base-pairing observables in ref. 28.

CJ signals showed variable agreement with annotated structures across RNA families (Fig. 1C, D). In tRNAs, CJ achieves high NormAUPR values and exceeds those of biophysical models. Intermediate behavior is observed in 16S and 5S rRNAs, where RNA-FM performs comparably to biophysical models, while Evo 2 and gLM2 show reduced performance. However, in all other RNA families, CJ signals underperform relative to biophysical models. We also analyzed 3,000 Rfam^30^ sequences spanning 11 RNA families (Fig. 1E), and found similarly that CJ-derived NormAUPR varies widely across families, whereas biophysical models remain comparatively more consistent. These results indicate that pairwise dependencies in gLMs can capture structure-associated signals, but the strength and reliability of their CJ signals vary substantially across RNA families.

A natural hypothesis based on the application of CJ to protein language models is that CJ signals reflect coevolutionary statistics^23^, where compensatory mutations between paired nucleotides can generate correlated sequence patterns detectable from sequence alignments. To test this possibility, we analyzed our Rfam-derived RNA dataset using R-scape^17^ to better understand to what extent covariation could account for CJ-predicted interactions within annotated structural pairs. Each base-pair was classified according to whether its CJ value passed a percentile-based cutoff (see Methods), whether it is predicted to have p(ij) ≥ 0.5 by EternaFold, and whether R-scape provides support for evolutionary covariation. Fig. 2A schematically illustrates the distinct types of pairing signals captured by R-scape and nearest-neighbor thermodynamic models such as EternaFold.

**Figure 2.**
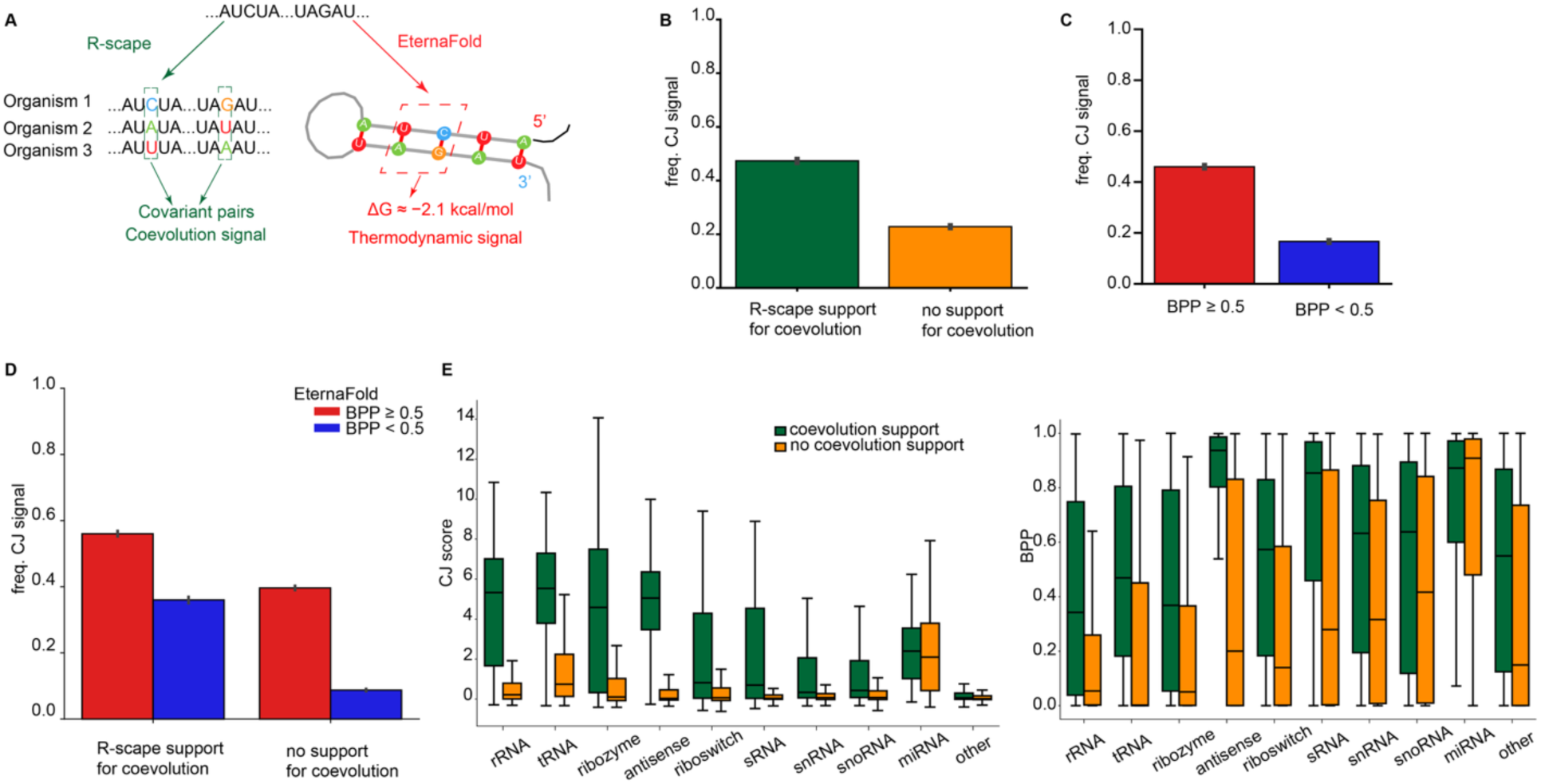
CJ signal associates with both evolutionary covariation and biophysical base-pairing probability. **(a)** We compared CJ signals to pairwise interactions predicted by R-scape and EternaFold: R-scape calculates evidence for coevolutionary support for base-pairs, whereas EternaFold performs a nearest-neighbor thermodynamic-based calculation. **(b)** Frequency of RNA-FM CJ-predicted base pairs in reference structures, stratified by R-scape covariation support. Base pairs supported by R-scape are more frequently predicted by CJ than pairs without detectable covariation. **(c)** Frequency of RNA-FM CJ-predicted base pairs in reference structures, stratified by EternaFold base-pairing probability (BPP > 0.5). Base pairs strongly supported by EternaFold are more frequently predicted by CJ signal. **(d)** When stratified by both R-scape coevolution support and EternaFold base-pairing probability (BPP > 0.5), CJ predicts substantial base-pairs with no R-scape coevolutionary support yet with EternaFold support. **(e)** Family-level comparisons of covarying and non-covarying base pairs for CJ and EternaFold. Both CJ and EternaFold assign stronger signals to covarying pairs.

We first examined CJ-prediction frequency separately for base pairs supported by R-scape and EternaFold. Base pairs supported by R-scape were more frequently predicted by CJ than those without detectable covariation, indicating that evolutionary covariation does indeed contribute to CJ signals (Fig. 2B). Similarly, base pairs with p(ij) ≥ 0.5 in the biophysics-based nearest-neighbor model EternaFold were more likely to be predicted by CJ than those with p(ij) < 0.5 (Fig. 2C). We next examined these signals jointly. Base-pairs supported by both R-scape and EternaFold showed the highest CJ-prediction frequency overall, whereas base-pairs lacking support from both methods showed the lowest frequency (Fig. 2D). Strikingly, we observed that many base pairs lacking detectable covariation but supported by EternaFold were still frequently predicted by CJ. These results indicate that covariation as detectable by R-scape alone accounts for only a small portion of CJ signals. Similar trends were also observed in Evo 2 and gLM2 (Supplementary Fig. 1A).

We observed that the degree to which the CJ signal from RNA-FM agreed with R-scape-supported base-pairs varied as a function of the RNA family (Fig. 2E). However, we also noticed something curious: EternaFold, the physics-based model, tended to have higher p(ij) for base-pairs with R-scape-supported coevolution than base-pairs without R-scape-supported coevolution (Fig. 2E, right). EternaFold’s training might confound this, as it was trained in part to maximize the likelihood of natural RNA structures, but the same trend holds for ViennaRNA which was fit entirely from biophysical measurements (Supplementary Fig. 1B).

### CJ correlates with biophysical base-pairing probabilities

Given our observation that some base pairs lacking R-scape support were supported by biophysical models, we next asked whether CJ-derived interactions show a systematic relationship to biophysically modeled base-pairing probabilities (BPPs). This question was also motivated by observations such as a GTP aptamer (bpRNA_PDB_245), in which we identified a helix with strong CJ signal that is absent from the annotated reference structure. Notably, this same helix is supported by elevated base-pairing probabilities in EternaFold, indicating that both CJ and biophysical models highlight a consistent interaction absent from the annotation (Fig. 3A). This helix appears in an alternative conformation predicted by EternaFold that is incompatible with the experimentally determined PDB structure (Fig. 3B). In the case of this GTP aptamer, the experimentally observed structure is likely stabilized by ligand binding, which could shift the structure ensemble away from a lower-energy conformation in the absence of ligand^31,32^. We were curious if this observation suggests that CJ-derived signals might reflect thermodynamic preferences not captured by single annotated structures but are instead consistent with alternative states in the free energy landscape.

**Figure 3.**
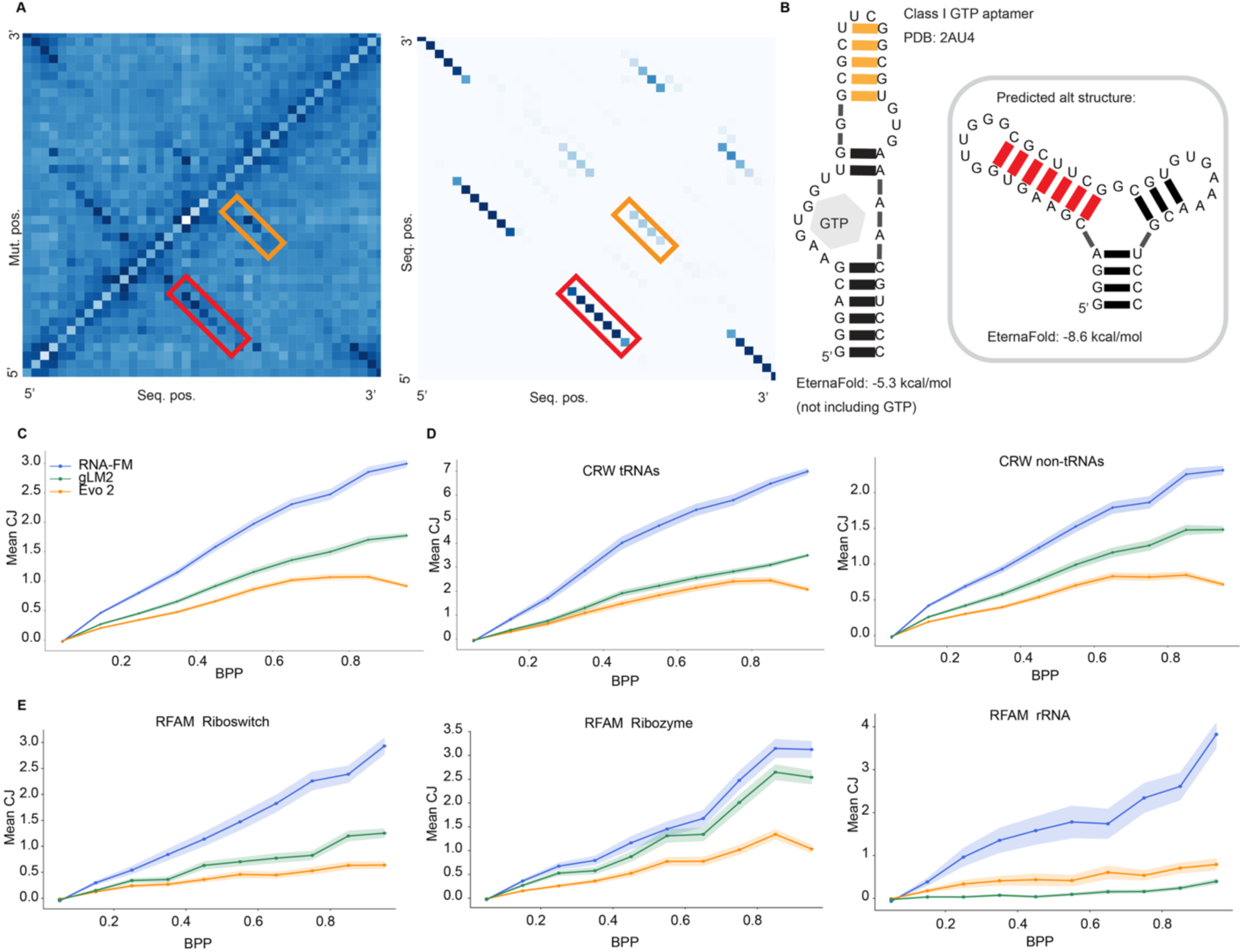
CJ values correlate with biophysical base-pairing probabilities. **(a)** CJ interaction map from RNA-FM (left) and EternaFold base-pairing probabilities (BPPs) (right) for a GTP aptamer (*bpRNA_PDB_245*). Base-pairing corresponding to annotated structural elements with weak CJ/BPP support are indicated in yellow. Alternative base-pairing supported by both CJ and BPPs, but absent from the annotated structure, is highlighted in red. **(b)** Secondary structures corresponding to these two states with relative free energies calculated in EternaFold. The alternative helix corresponds to a thermodynamically favorable state that may not be realized under ligand-bound conditions. **(c)** Relationship between CJ values and BPPs predicted by EternaFold across all nucleotide pairs. Nucleotide pairs show an increase in CJ signal with increasing pairing probability across gLMs. The same trend is true in **(d)** tRNA and non-tRNA sequences within the CRW dataset **(e)** families in the Rfam dataset with sufficient representative members.

To systematically test this, we evaluated the correlation between EternaFold BPPs and CJ values, and to our surprise, found that BPPs trended consistently with CJ values (Fig. 3C). This relationship is observed across all three gLMs, although the magnitude of the trend varies between models, with the highest being RNA-FM and the lowest being Evo 2 (Pearson r: 0.53, 0.56, and 0.43 for RNA-FM, gLM2, and Evo 2, respectively). We were curious if this trend is driven by families predicted best by CJ, and instead found the trend held across several families, including both tRNA and non-tRNA subsets in the CRW dataset (Fig. 3D) and several families in Rfam (Fig. 3E).

### “Mirror test” preserves biophysical structure prediction but disrupts CJ-derived pairing signals

The positive association between CJ values and biophysical base-pairing probabilities (Fig. 3) raises the possibility that CJ-derived signals reflect dependencies like those captured by biophysical folding models. If CJ-derived signals primarily reflect thermodynamics, they should be robust to transformations that preserve thermodynamics. We designed a simple “mirror test” to evaluate this: each RNA sequence was reversed from 5′→3′ to 3′→5′ and used as input into biophysical models or the CJ framework (Fig. 4A). This transformation preserves sequence and structure content, and under which base-pairing relationships remain relatively stable in biophysical models (Fig. 4B,C). These reversed structures should not be taken as a ground truth for the reversed sequence, given the prevalence of cotranscriptional folding in nature, but offer a means to compare nearest-neighbor-based structure predictions to CJ signals.

**Figure 4.**
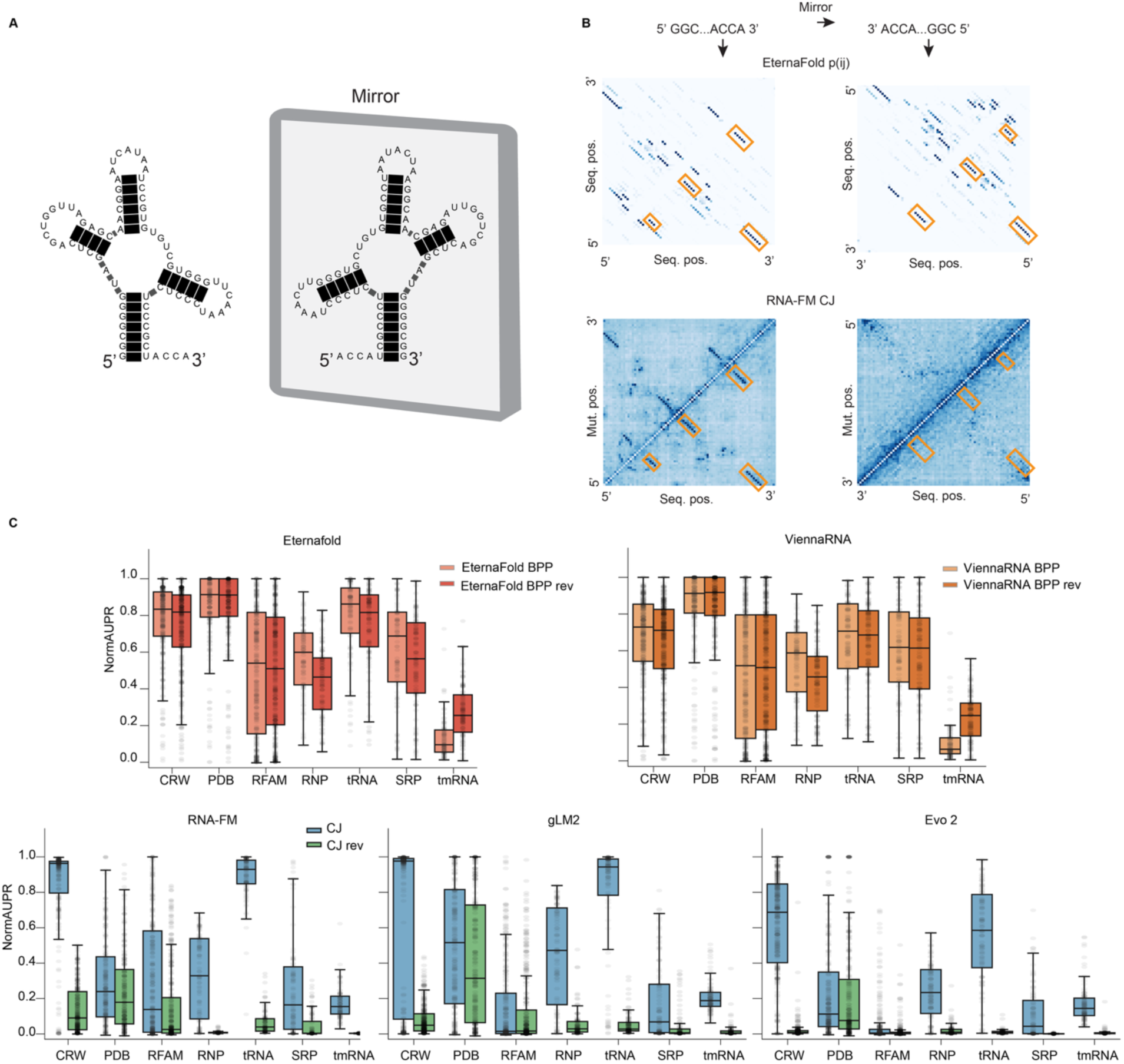
“Mirror test” preserves biophysical structure prediction but disrupts CJ-derived pairing signals. **(a)** Schematic illustration of the “mirror test”: an RNA sequence is reversed from its original 5’→3’ orientation to 3’→5’, and the corresponding annotated secondary structure is also reversed. **(b)** Comparing RNA-FM CJ maps and EternaFold BPP maps before and after mirror reversal for a tRNA (bpRNA_CRW_30836). EternaFold predictions remain qualitatively similar after reversal, whereas RNA-FM CJ signals are substantially disrupted. **(c)** NormAUPR values for original and mirror-reversed sequences across gLMs and biophysical models. Biophysical models retain largely similar performance after reversal, whereas CJ-derived signals from all three gLMs are strongly reduced toward baseline.

For biophysical models (EternaFold and ViennaRNA), mirror reversal results in largely similar NormAUPR values compared to the original sequences, indicating that predicted base-pairing probabilities are relatively robust to sequence reversal (Fig. 4C). In contrast, CJ-derived signals from all three gLMs are strongly disrupted by mirror reversal. Regardless of their original performance, NormAUPR values decrease to nearly zero after reversal, indicating a loss of structure-associated signal. This effect is illustrated in the same representative tRNA example (bpRNA_CRW_30836) shown in Fig. 1, where EternaFold predictions remain qualitatively consistent after reversal, while CJ signals from RNA-FM are substantially degraded (Fig. 4B). These results indicate that CJ-derived signals are not invariant under sequence reversal and instead depend strongly on relative nucleotide positioning in the input sequence.

### Thermodynamics in CJ can confound interpretation of evolutionary conservation, but synthetic controls offer a path forward

Recent work has interpreted CJ-derived pairwise dependencies in RNA language models as evidence of learned evolutionary or co-evolutionary constraints, including in the preQ1 riboswitch system^10^. However, our earlier analyses suggest that CJ signals may also emerge from thermodynamic or broader statistical regularities independent of evolutionary conservation. These observations motivate the need for additional controls when interpreting CJ-derived interactions as evidence of conserved RNA structure.

We started by examining a simple structural element of a long noncoding RNA whose structural conservation has been debated, helix 11 in MEG3 (structure in Fig. 5A, top). Previous work proposed helix 11 is evolutionarily conserved^16^, while subsequent analyses argued that this conclusion may arise from inappropriate statistical interpretation of covariation signals ^19^(Fig. 5B, top). We evaluated helix 11 using RNA-FM CJ and observed weak but detectable interaction signals in the region corresponding to the proposed stem. However, to determine whether such signals necessarily reflect evolutionary constraint, we constructed a set of random RNA sequences that are predicted by EternaFold to adopt the same secondary structure. Notably, these synthetic sequences also produce CJ interaction patterns with stems in the same location, despite lacking any evolutionary relationship (Fig. 5C, top). This demonstrates that CJ-derived interactions can arise purely from intrinsic thermodynamic stability, independent of evolutionary covariation.

**Figure 5.**
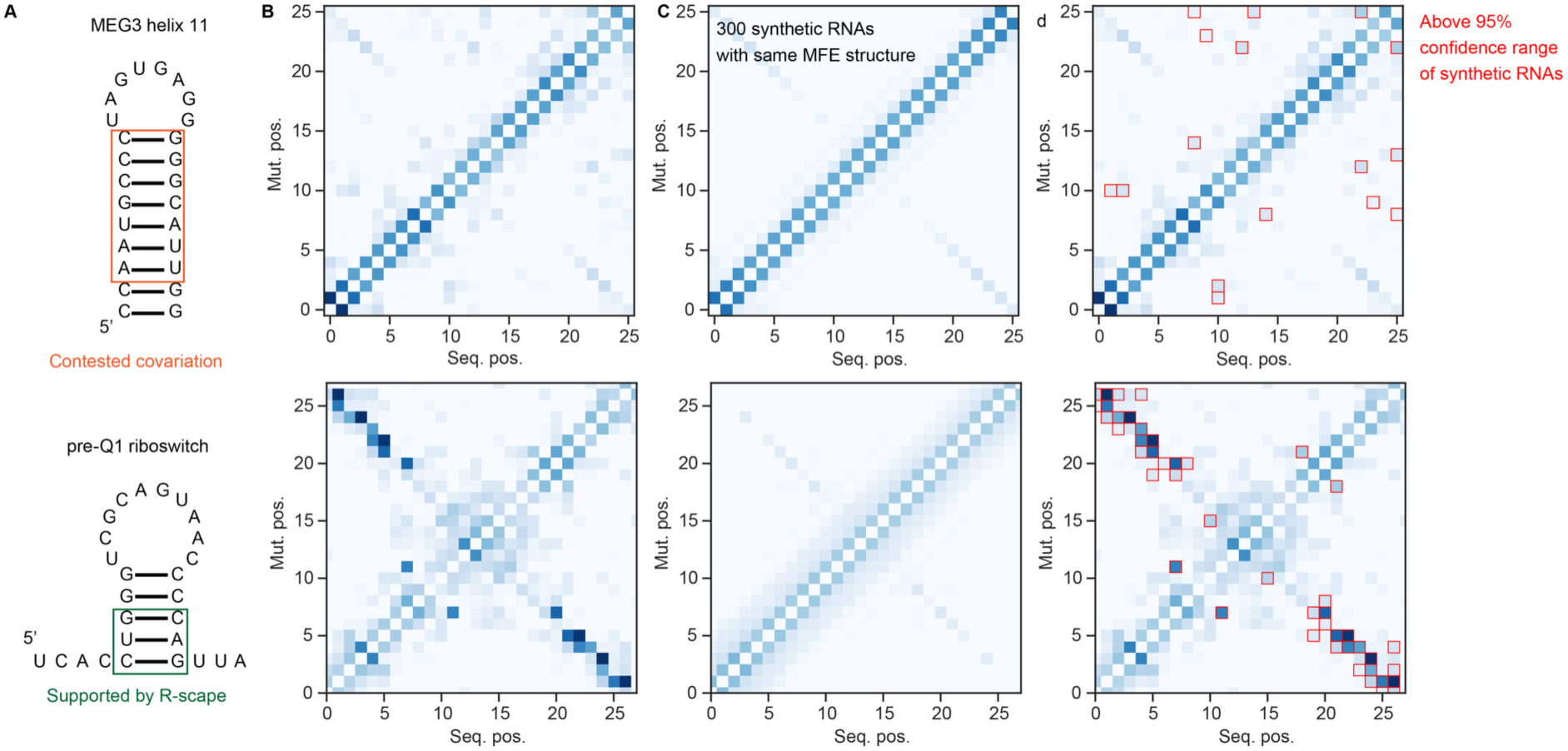
Comparing CJ signals to those from synthetic control RNAs reveals signals not consistent with “thermodynamic” baseline. **(a)** Secondary structures of MEG3 helix 11 (top), whose covariation is contested, and the pre-Q1 riboswitch (bottom), whose covariation is supported by R-scape. Positions supported or contested by R-scape are boxed in green and orange, respectively. **(b)** CJ heatmaps for each RNA. Both MEG3 helix 11 and the pre-Q1 riboswitch display CJ signals at base-pairing positions. **(c)** Mean CJ heatmaps computed from 300 synthetic RNA sequences generated to fold to the same minimum free energy (MFE) structure as each natural sequences, serving as a structure-matched null distribution. Notably, there is signal at the base-pairing positions in both sets of controls. **(d)** Positions where the observed CJ values (b) exceed the 95^th^ percentile of the synthetic null distribution (c) are boxed in red. In MEG3 helix 11, the above-null positions are scattered and not centered on known base-pairing regions, suggesting that the observed covariation pattern as shown by CJ is not distinguishable from that learned from structural patterns alone. In contrast, in the pre-Q1 riboswitch, the signal above the null is concentrated around known base-paired positions.

To move beyond this confounding factor, we propose a significance test that explicitly accounts for structure-derived signals by treating the distribution of CJ contacts across structure-matched synthetic sequences as a null. Positions where the observed CJ signal exceeds the 95th percentile of this null are candidates for evolutionarily meaningful constraint, distinct from what “learned thermodynamics” alone would predict. Applying this framework to the MEG3 helix 11 demonstrates that positions above the 95^th^ percentile are diffuse and not in the designed synthetic base-paired regions that show signal in the control distribution (Fig. 5D, top).

Applying this same framework to the pre-Q1 riboswitch, whose covariation is well supported by R-scape, reveals above-null signal concentrated at known base-pairing positions (Fig. 5A-D, bottom), as well as other pairwise dependencies. This approach allows for evaluating model-derived pairwise dependencies that are distinct from those learned from base-pairing alone and may be more broadly useful for making claims about novel patterns detected by gLMs. We note that this significance test would not be required if a postulated structure shows no signal in CJ: we generated RNA-FM CJs for the long non-coding RNA HOTAIR, whose evolutionary conservation has been contested, and found no evidence of CJ signal for it across three organisms (Supplementary Figure 2).

## Discussion

In this work, we investigated what categorical Jacobian (CJ) signals capture in genomic language models (gLMs) and how evolutionary covariation vs. thermodynamics might account for signals related to RNA structure. We first found that the predictive power of CJ-derived signals varies substantially across RNA families. This indicates that gLMs capture structure-associated dependencies, but that these signals are not expressed uniformly across sequence contexts. One hypothesis for how gLMs encode RNA structure is that they might have learned a route to compress statistics for every RNA family in the training set, akin to what has been postulated for protein language models and perhaps in something like a context-free grammar, as has been postulated in ref. 23. However, the mixed performance of CJ signals by family discredits this hypothesis. For gLM-based models that perform supervised training from language model embeddings to predict RNA structure, this suggests that the supervision process is more important than it might be in the protein context. Similar observations were reported by Teng et al^33^.

We queried to what extent evolutionary covariation detected by the standard method R-scape could account for structure-associated CJ signals and found surprisingly small overlap between R-scape and CJ signals (Fig. 2B). Evolutionary covariation does contribute to CJ enrichment within annotated base-pairs, similarly to findings originally described for protein language models^23^, but it does not fully account for the observed signal. Doubtlessly some CJ-predicted base pairs that lack R-scape support would have R-scape support if deeper multiple sequence alignments were available. Notably, CJ predicts a substantial fraction of base pairs with no R-scape support, yet which are predicted by biophysical base-pairing probabilities from the physics-based model EternaFold.

We designed a “mirror test” to probe the extent to which gLMs reflect RNA thermodynamics for sequences that are out-of-distribution. This test involves reversing natural RNA sequences 3’->5’ and mirroring their reference structures, which creates a matched set of controls that preserve sequence and local structure content while removing the possibility of a model using learned pairwise covariation or other patterns. A model that understands RNA folding grammar, such as nearest-neighbor models, should capture similar propensities for the reference structure, given that local energetics are comparable. However, we found that gLMs suffered in this mirror test, underscoring that gLM pairwise associations do not behave like nearest-neighbor models.

Considering the failure of the gLMs to pass the “mirror test”, the extent to which they can capture some thermodynamic properties is striking, even to the extent that their CJ values trend with biophysically predicted base-pairing probability values. This demonstrates the capacity of gLMs to “learn” patterns of natural RNA sequences that appear to be consistent with nontrivial biophysical properties like thermodynamics. It is tempting to speculate that this might be related to our finding that base pairs with R-scape support have higher predicted base-pairing values: evolutionary covarying base-pairs could have sufficient information to “learn” thermodynamics.

One possible concern with CJ is that perturbing the input sequence and monitoring output changes may not necessarily reflect the internal representations learned by the model itself for nucleotides. Recently, Teng et al. introduced REDIAL^33^, a related perturbation-based framework that instead probes perturbation responses directly from intermediate embedding representations. Despite these methodological differences, we found that applying REDIAL to our key analyses reproduced our same conclusions, including relationships between pairwise signals, evolutionary covariation, and thermodynamic pairing propensity (Supplementary Fig. 3). These observations suggest that the major conclusions described here are not specific to a single perturbation-based interpretation method but instead reflect broader properties of RNA language model representations.

If a gLM detects a pairwise dependency that is not supported by covariation analysis, does this count as a false positive? Or could the signal be a conserved evolutionary pattern beyond the scope of independent pairwise conservation patterns that current standard covariation analysis detects? Our MEG3 case study shows that synthetic RNAs with designed thermodynamics can generate CJ signal, and therefore any claim of evolutionary conservation requires more careful interpretation. The synthetic control framework we introduce here offers a route to distinguish significant pairwise signals in a gLM. By generating sequence-diverse, structure-matched synthetic sequences and identifying CJ signals that exceed the resulting distribution, one can isolate positions where a gLM infers signal beyond what thermodynamics alone would predict. This framework makes no assumption about what that additional signal represents: it could reflect genuine evolutionary covariation, higher-order structural context, or interactions with the broader sequence environment.

We note that constructing well-matched synthetic controls is straightforward for simple helix elements, where random sequences constrained to a target structure can be generated efficiently and folding predictions are reliable. For more complex RNA architectures, including elements with multiple junctions, pseudoknots, or tertiary contacts, generating truly de novo structure-matched nulls becomes more challenging, but not insurmountable; tools from the RNA design community, such as NEMO^34^, EternaBrain^35^, and others, are capable of generating synthetic sequences that fold to target structures under thermodynamic constraints, and would be directly applicable to constructing control distributions for more complex elements.

An important next step in improving gLMs will be to create architectures that can explicitly separate contributions from evolution and thermodynamics. More refined benchmarks, especially those incorporating synthetic controls, may help distinguish structure-relevant signals from broader sequence statistics. In this work, we only analyzed models at a sequence-only level of training, but models that incorporate high-throughput experimental observables in training offer richer paths for understanding how sequence- and observable-based training intersects^36^. More generally, using the principles described here to understand how model behavior changes with different training architectures and training datasets, as well as across RNA families, sequence lengths, and compositional regimes, will be essential for building more interpretable gLMs and for best using them in biomolecular discovery and design.

## Acknowledgments

We thank Christian Macdonald, Gina El Nesr, Rhiju Das, Shujun He, and members of the Wayment-Steele lab for useful discussion. This work was supported by startup funds from the University of Wisconsin-Madison School Department of Biochemistry.

## Author contributions

YX, NNP, and HKWS designed the research and contributed to data analysis. YX and HKWS wrote the manuscript. All authors provided manuscript feedback.

## Data/code availability

Representative code to reproduce the findings presented here is available at https://github.com/WaymentSteeleLab/Evo_or_Thermo

## AI usage statement

Claude and ChatGPT were used for light editing assistance.

## Methods

### RNA datasets and structure annotations

We obtained RNA sequences and their corresponding annotated secondary structures from the bpRNA-1m(90)^29^ dataset. This resource aggregates RNA sequences from multiple sources, including CRW^37^, PDB^38^, SRP^39^, SPR (tRNA-only)^40^, tmRNA^41^, RNP^42^, and Rfam^30^. To construct the main evaluation set, we randomly sampled sequences from seven source datasets: 300 sequences each from CRW and Rfam, 250 from PDB, and 100 each from SRP, SPR, tmRNA, and RNP, yielding a total of 1,250 sequences. We excluded sequences with ≥1,024 nucleotides due to model input constraints. For the analyses in Fig. 1 and Fig. 2, we used six non-Rfam source datasets, comprising 950 sequences in total. The originally sampled 300 Rfam sequences were not used for these family-level analyses; instead, we generated an expanded Rfam dataset to better assess family-specific variation. Specifically, we randomly sampled 3,000 RNA sequences from the Rfam subset of bpRNA-1m(90), which were used for the Rfam family-stratified analyses in Fig. 1 and Fig. 2.

For the mirror test in Fig. 4, we used the original set of 1,250 sequences, consisting of 950 sequences from the six non-Rfam source datasets together with the originally sampled 300 Rfam sequences. This ensured that the mirror test benchmark retained equal representation across the seven bpRNA source datasets.

### Genomic language models and biophysical models

We evaluated three genomic language models trained on distinct sequence corpora: RNA-FM, which is trained on RNA sequences; Evo 2-1b, trained primarily on genomic DNA sequences; and gLM2-650M, trained on metagenomic data. These models were selected to represent a range of training data distributions. For all models, we used the publicly available pretrained weights without any task-specific fine-tuning.

To compare with nearest-neighbor biophysical models, we used EternaFold (v1.3.1) and ViennaRNA RNAfold (v2.7.2) to compute base-pairing probability (BPP) matrices. BPP matrices, free energies, and Boltzmann-sampled substates (cf Fig. 3B) for both EternaFold and ViennaRNA were calculated using default settings in the arnie^43^ package. ViennaRNA calculations were performed with the temperature parameter set to 60°C, as shown in ref. 28 to improve performance over default temperature. Unless otherwise specified, default parameters were used for both biophysical models.

### Categorical Jacobian (CJ) computation

We followed the CJ computation described in ref. 23. We briefly summarize here as follows: for an input RNA sequence *x* of length *L*, let *f_j_*(*x*)*_b_* denote the model logit at position *j* for nucleotide *b*. For each position *i*, we generated perturbed sequences *x*^(*i→a*)^ by substituting the nucleotide at position *i* with nucleotide *a* ∈ {*A, C, G, U*}, and computed the resulting change in logits:

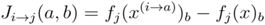

This yields a four-dimensional tensor *J* ∈ ℝ^*L*×4×*L*×4^, representing the sensitivity of outputs at position *j* to perturbations at position *i*. To reduce background bias, we applied sequential mean-centering along each tensor dimension, subtracting the mean across each axis in turn. A scalar interaction score between positions *i* and *j* was then computed by taking the Frobenius norm over the categorical dimensions:

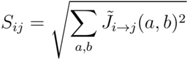

The resulting matrix was symmetrized as:

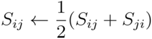

and diagonal entries were set to zero.

To further remove global biases, average product correction (APC) was applied:

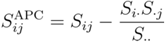

where *S_i._*, *S_.j_*, and *S*. denote row sums, column sums, and the total sum of the matrix, respectively.

### Evaluation against RNA secondary structure (NormAUPR)

To evaluate the extent to which CJ-derived signals recover annotated RNA secondary structure, we compared model-derived interaction scores to annotated base pairs using the area under the precision–recall curve (AUPR). For each RNA sequence of length L, we constructed a binary label vector indicating whether each nucleotide pair (i, j) corresponds to an annotated base pair. The corresponding prediction scores were taken from the CJ-derived interaction matrix (or BPP matrix for biophysical models). All nucleotide pairs were ranked according to their interaction scores, and precision–recall curves were computed by progressively including pairs in descending order of score. AUPR was calculated using a step-wise integration of the precision–recall curve.

Because the proportion of true base pairs varies across RNA sequences, raw AUPR values are not directly comparable across sequences of different lengths or structural densities. To account for this, we computed a normalized AUPR (NormAUPR) for each sequence by adjusting for the baseline AUPR expected under random ranking. Specifically, the baseline was defined as the fraction of positive pairs among all possible nucleotide pairs. The normalized AUPR was then computed as:

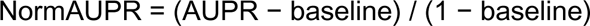

This normalization maps random performance to 0 and perfect recovery of annotated base pairs to 1, enabling comparison across sequences and datasets. Unless otherwise specified, AUPR and NormAUPR were computed independently for each sequence and then aggregated for downstream analyses.

### Covariation analysis using R-scape

To assess the relationship between CJ-derived signals and evolutionary covariation, we performed covariation analysis using R-scape on RNA sequences from the Rfam subset of bpRNA-1m(90). The same set of 3,000 Rfam sequences used for the family-level analyses in Fig. 1 and Fig. 2 was used throughout. Multiple sequence alignments (MSAs) for each RNA family were obtained from the corresponding Rfam entries in Stockholm (.sto) format. These MSAs were used as input to R-scape to identify nucleotide pairs with significant covariation. For all analyses in this study, we used the structure-informed two-set R-scape analysis. During analysis, only annotated base pairs from the reference secondary structure were considered and each annotated pair was classified as either covarying or non-covarying based on R-scape output. Annotated secondary structures were obtained from the corresponding Rfam annotations.

For the analyses in Fig. 2B-D, we considered only nucleotide pairs (i,j) annotated as base pairs in the reference secondary structure. Within this set, each pair was classified according to three binary criteria: whether it was predicted by CJ, whether its EternaFold base-pairing probability exceeded 0.5, and whether it was identified as covarying by R-scape. A nucleotide pair was defined as CJ-predicted if its CJ value ranked within the top 3% of values in row i, within the top 3% in column j, and within the top 3% of all pairwise CJ values for that sequence. To focus on non-local interactions, nucleotide pairs with |i-j| <= 6 were excluded from this classification. We then computed the frequency of CJ-predicted pairs within groups defined by R-scape and EternaFold support.

### Correlation analysis

For the correlation analysis in Fig. 3C-E, we quantified the relationship between CJ signal strength and biophysical base-pairing probabilities across all base pairs. For each base pair, the corresponding CJ interaction value and EternaFold BPP were retrieved from the CJ matrix and BPP matrix, respectively.

To analyze how CJ signal varies as a function of thermodynamic support, base pairs were grouped according to their EternaFold BPP values using equally spaced BPP bins. Within each BPP bin, we collected the corresponding CJ values and computed the mean CJ signal across all nucleotide pairs assigned to that bin. Confidence intervals were estimated independently for each bin using the standard error of the mean:

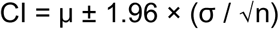

where μ denotes the mean CJ value within the bin, σ is the sample standard deviation, and n is the number of nucleotide pairs in the bin. Confidence intervals therefore approximate the 95% confidence interval under a normal approximation.

This analysis was performed independently for RNA-FM, gLM2, and Evo 2. Mean CJ values and their corresponding confidence intervals were visualized as line plots across BPP bins to assess the extent to which stronger thermodynamic base-pairing support was associated with increased CJ signal across models.

### Mirror test

To assess the dependence of model-derived signals on sequence orientation, we performed a mirror test by reversing each RNA sequence from its original 5′→3′ orientation to 3′→5′. For each sequence, a reversed version was generated by inverting the nucleotide order. The corresponding annotated secondary structure was transformed accordingly by reversing nucleotide indices, ensuring that base-pairing relationships were preserved under the reversal.

Both the original and reversed sequences were used as inputs to gLMs and biophysical models. For each case, CJ-derived interaction matrices and BPP matrices were computed as described above. The ability of these signals to recover annotated base pairs was then evaluated using the same AUPR and NormAUPR metrics.

### Significance test using synthetic sequences

To construct random sequences, we randomly sampled base-pairs and unpaired nucleotides and calculated the MFE structure of the resulting sequence in EternaFold. If the MFE structure matched the target structure, the sequence was kept. CJs were calculated for these sequences using the same protocol in RNA-FM as described previously. A pairwise dependence i:j of a natural construct was deemed significant (boxed interactions in red in Fig. 5) if CJ at position i:j was greater than the 97.5% percentile (i.e., above 95% confidence interval) of 300 sampled random sequences for each natural construct.

---

**Supplementary Figure 1.**
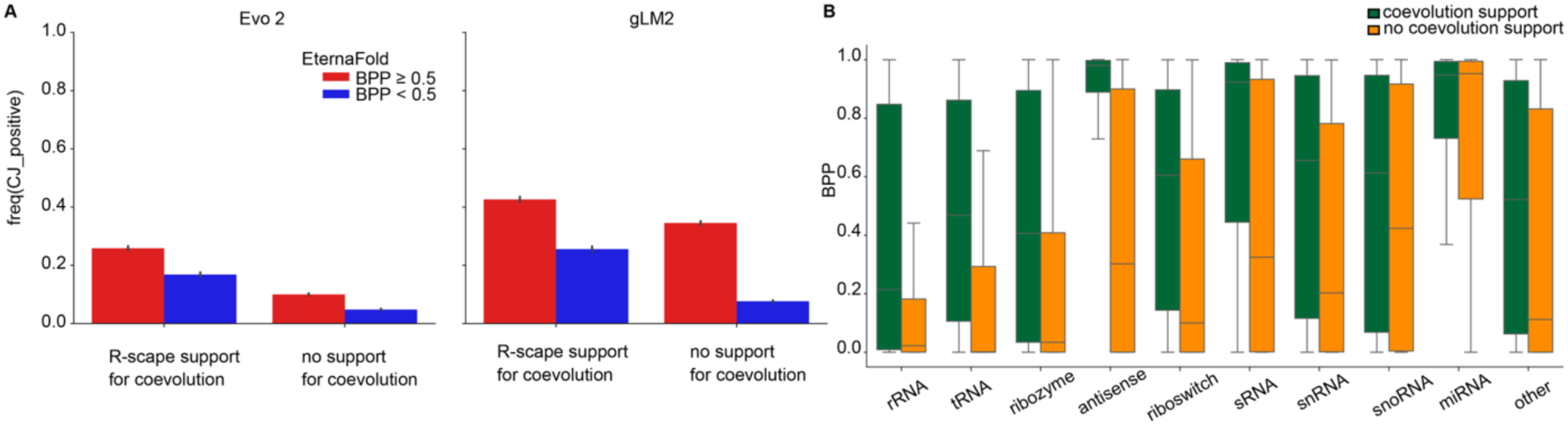
CJ signal is explained by both evolutionary covariation and biophysical base-pairing probability across multiple models. **(a)** Frequency of CJ-predicted base pairs among base pairs in reference structures, stratified by R-scape support and EternaFold base-pairing probability (BPP > 0.5), for Evo 2 and gLM2 (compare to Fig. 2D). **(b)** Family-level comparisons of covarying and non-covarying base pairs for ViennaRNA (compare to Fig. 2E).

**Supplementary Figure 2.**
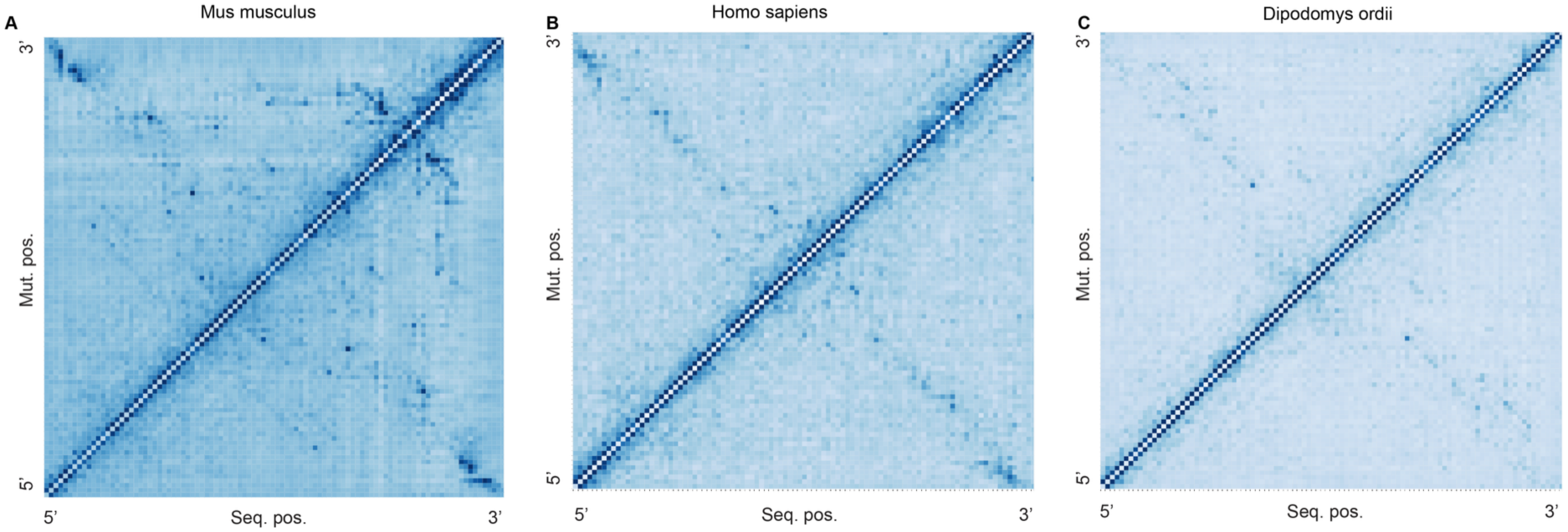
Variation of CJ-derived structure-associated signals across HOTAIR_3 homologs. **(a-c)** CJ interaction maps for three HOTAIR homologous sequences from different species: (a) AAHY01125231.1 (*Mus musculus*), (b) DQ926657.1 (*Homo sapiens*), and (c) ABRO01112174.1 (*Dipodomys ordii*). While all sequences correspond to the same annotated HOTAIR_3 region, the strength CJ signals vary substantially across homologs. This variability suggests that CJ-derived structural signal is not uniformly preserved across species, highlighting that apparent model support does not necessarily reflect a stable or conserved structural constraint.

**Supplementary Figure 3.**
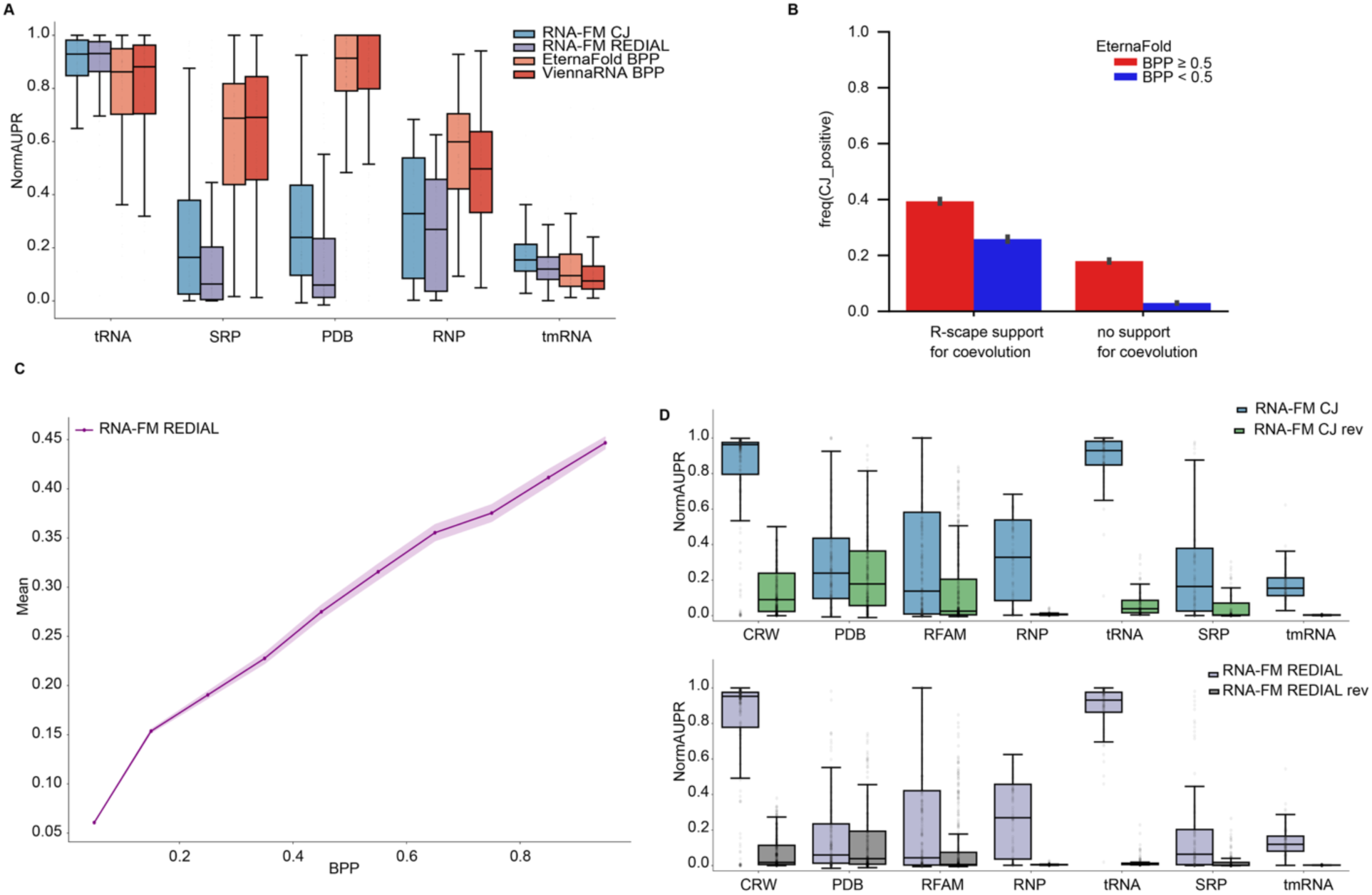
REDIAL recapitulates major trends observed with CJ-derived signals. **(a)** Distribution of normalized area under the precision–recall curve (NormAUPR) for RNA-FM CJ and REDIAL across five RNA datasets, quantifying agreement between model-derived signals and annotated secondary structures (compare to Fig. 1C). **(b)** Frequency of REDIAL-predicted base pairs in reference structures, stratified by R-scape coevolution support and EternaFold base-pairing probability (BPP > 0.5). Similar to CJ-derived signals, base pairs supported by both evolutionary covariation and thermodynamic pairing propensity show the highest prediction frequency (compare to Fig. 2D). **(c)** Relationship between REDIAL values and EternaFold-predicted BPPs across all nucleotide pairs. Similar to CJ-derived signals, REDIAL values increase with increasing thermodynamic pairing probability (compare to Fig. 3C). **(d)** NormAUPR values for original and mirror-reversed sequences using REDIAL-derived signals. Similar to CJ-derived signals, REDIAL signals are strongly reduced after sequence reversal, whereas biophysical nearest-neighbor models remain largely unchanged (compare to Fig. 4C).

